# Diagnostic Yield and Treatment Impact of Targeted Exome Sequencing in Early-onset Epilepsy

**DOI:** 10.1101/139329

**Authors:** Michelle Demos, Ilaria Guella, Marna B. McKenzie, Sarah E. Buerki, Daniel M. Evans, Eric B. Toyota, Cyrus Boelman, Linda L. Huh, Anita Datta, Aspasia Michoulas, Kathryn Selby, Bruce H. Bjornson, Gabriella Horvath, Elena Lopez-Rangel, Clara DM van Karnebeek, Ramona Salvarinova, Erin Slade, Patrice Eydoux, Shelin Adam, Margot. I. Van Allen, Tanya N. Nelson, Corneliu Bolbocean, Mary B. Connolly, Matthew J. Farrer

## Abstract

**Background:** To examine the impact on diagnosis, treatment and cost with early use of targeted whole-exome sequencing (WES) in early-onset epilepsy.

**Methods:** WES was performed on 50 patients with early-onset epilepsy (≤ 5 years) of unknown cause. Patients were classified as retrospective (epilepsy diagnosis > 6 months) or prospective (epilepsy diagnosis < 6 months). WES was performed on an Ion ProtonTM and variant reporting was restricted to the sequences of 565 known epilepsy genes. Diagnostic yield and time to diagnosis were calculated. An analysis of cost and impact on treatment was also performed.

**Results:** A likely/definite diagnosis was made in 17/50 patients (34%) with immediate treatment implications in 8/17 (47%). A possible diagnosis was identified in 9 additional patients (18%) for whom supporting evidence is pending. Time from epilepsy onset to genetic diagnosis was faster when WES was performed early in the diagnostic process (mean: 143 days prospective versus 2,172 days retrospective). Costs of prior negative tests averaged $8,344 in the retrospective group, suggesting savings of up to $5,110 per patient.

**Interpretation:** These results support the clinical utility and potential cost-effectiveness of using targeted WES early in the diagnostic workup of patients with unexplained early-onset epilepsy. The costs and clinical benefits are likely to continue to improve. Advances in precision medicine and further studies regarding impact on long-term clinical outcome will be important.

## Introduction

Epilepsy is a common pediatric neurological disorder with increased risk of developmental delay, autism and psychiatric illness, for which treatment is ineffective in 30-40% of patients. High-throughput sequencing technologies, including whole-exome sequencing (WES) and epilepsy gene panels, have advanced our genetic understanding as pathogenic variants have been identified in 10-78% of select patients (1–6). A genetic diagnosis of epilepsy may enable more accurate counseling regarding prognosis and recurrence risk, avoids unnecessary medical investigations and may change care. It also allows families to connect with the same genetic condition and/or join support groups. Recent studies have demonstrated the potential cost savings of WES in the diagnostic work-up of children with suspected monogenic disorders (7–10). However, in Canada, access to such technology in clinical care is variable. In this British Columbia study, we assess the effectiveness of using WES by comparing diagnostic yield, time to diagnosis, and cost to current clinical practices. The potential treatment impact of a genetic diagnosis is also described.

## Methods

### Patients

Fifty patients with epilepsy (11) were enrolled between December 2014 and June 2015. All had seizure onset at ≤5 years of undefined cause after EEG, brain MRI and chromosome microarray investigations. Seizure types and electroclinical syndromes were classified according to the International League Against Epilepsy (ILAE)(12). Patients with self-limiting benign electroclinical syndromes, such as Childhood Absence Epilepsy (onset >4 years), were excluded as they most likely have multifactorial inheritance. Patients were classified as **retrospective** (n=37), defined as an epilepsy diagnosis >6 months before study enrollment with a standard clinical approach to genetic testing (variable genetic tests which include gene-by-gene approach using Sanger sequencing (n=15), small epilepsy gene panels using high-throughput sequencing (n=4), and/or mitochondrial DNA sequencing (n=4)); or **prospective** (n=13), which included an epilepsy diagnosis <6 months before study enrollment date and having limited to no genetic testing. Varying degrees of screening tests for inborn errors of metabolism; such as plasma amino acids, lactate and ammonia, was also performed in both groups. Clinical data was recorded using a secure Research Electronic Data Capture (REDCap)(13) information system hosted at Child and Family Research Institute.

This study was approved by the BC Children’s Hospital and University of British Columbia Ethics Board. Informed consent and/or assent were obtained before study inclusion.

### Whole-exome sequencing

Genomic DNA was extracted from peripheral blood lymphocytes following standard protocols. Exonic regions were captured using the Ion AmpliSeq Exome Kit (57.7Mb) and WES was performed on an Ion Proton^TM^ according to manufacturers’ recommendations (Life Technologies Inc., CA) within 2 weeks of receiving samples. Reads were aligned against the human reference genome hg19. Variant annotation was performed with ANNOVAR (14) integrating data from PHAST PhyloP (15), SIFT (16), Polyphen2 (17), LRT (18) and MutationTaster (19) algorithms, Combined Annotation Dependent Depletion (CADD) scores (20), dbSNP (www.ncbi.nlm.nih.gov/SNP/), the Exome Aggregation Consortium (ExAC; exac.broadinstitute.org) and ClinVar (21)(www.ncbi.nlm.nih.gov/clinvar). Additionally, variants were compared to an in-house database containing more than 900 exomes to exclude platform artifacts and common variants not present in public databases.

Analysis was restricted to 565 genes previously implicated in epilepsy (Supplementary Table 1), using a gene-reporting pipeline developed in-house. The gene list was compiled through the combination of a comprehensive literature search (Pubmed, OMIM) and clinically available epilepsy panels (GeneDx, Courtagen). Annotation was limited to exonic nonsynonymous and splicing (±3bp) substitutions. Homozygous variants, potential compound heterozygous variants (defined as genes with >1 variant locus per individual) with a minor allele frequency (MAF) <5% and heterozygous variants with MAF <0.1% were reported. All samples were required to meet minimum quality standards, with a WES average coverage >80X.

Sanger sequencing, performed as previously described (22), was used on a case-specific basis in a few individuals with very specific clinical phenotypes to complete regions of poor coverage in genes related to the patient’s phenotype when no candidate variants were identified, or when a heterozygous and potentially pathogenic variant was identified in gene previously implicated in autosomal recessive disease. No additional variants were identified though post-WES Sanger sequencing.

### Variant prioritization and validation

Cases were reviewed at a bi-weekly meeting by a multi-disciplinary genomic team. Variant prioritization was performed based on: 1) frequency in public databases; 2) predicted protein impact; 3) disease inheritance, and; 4) correlation of patient phenotype and candidate gene literature. Up to 3 putative causative variants were validated by Sanger sequencing in patient and parental samples. Clinical Sanger sequencing confirmation and interpretation in accordance with ACMG guidelines (23) allowed disclosure to families and management adjustments when indicated. Two time intervals were measured: 1) from a clinical diagnosis of epilepsy to Sanger validation of a putative pathogenic variant; and 2) from enrollment with genetic counselling to Sanger validation of same.

### Genetic Counseling and Treatment Implications

Pre- and post-test genetic counseling was performed for each patient/family. As only a limited set of 565 genes related to seizure disorders were annotated, and only in affected probands, reporting related to incidental (secondary) findings was uncommon (24). Genetic disorders with specific therapeutic implications (47 genes) were defined as conditions in which current literature supports a preferred antiepileptic medication and/or approach (25–27).

### Cost Estimation

Resource use data were retrospectively acquired from electronic health records and medical charts. Cost estimates in Canadian dollars were based on micro-cost information from the British Columbia Provincial Medical Service Plan Index (2015), Canadian Interprovincial Reciprocal Billing Rates (2014/2015), Children’s and Women’s Health Centre of British Columbia Internal Fee Schedule (2015) and the internal accounting system. Diagnostic costs included: biochemical tests, imaging tests, genetic tests, neurophysiological tests and biopsies (a complete list of tests is provided in *Appendix 1*). Academic and/or hospital pricing is used throughout. Inpatient hospitalization costs, outpatient visits such as clinic visits, and indirect costs such as parental time off work for medical visits related to their child’s epilepsy were not included.

### Data Analysis

All categorical and quantitative variables were analyzed using STATA (Release 13, College Station, TX).

## Results

Targeted WES was performed on 50 patients and clinical features are summarized (Supplementary Table 2). The average age of epilepsy onset was 19 months (range 0.2-60 months), 18 months for prospective cases (n=13) and 19 months for retrospective cases (n=37)(Table 1). Of the 565 genes, 90% had at least 85% of their consensus-coding region sequenced with >20X coverage (Supplementary Table 1).

**Table 1.**
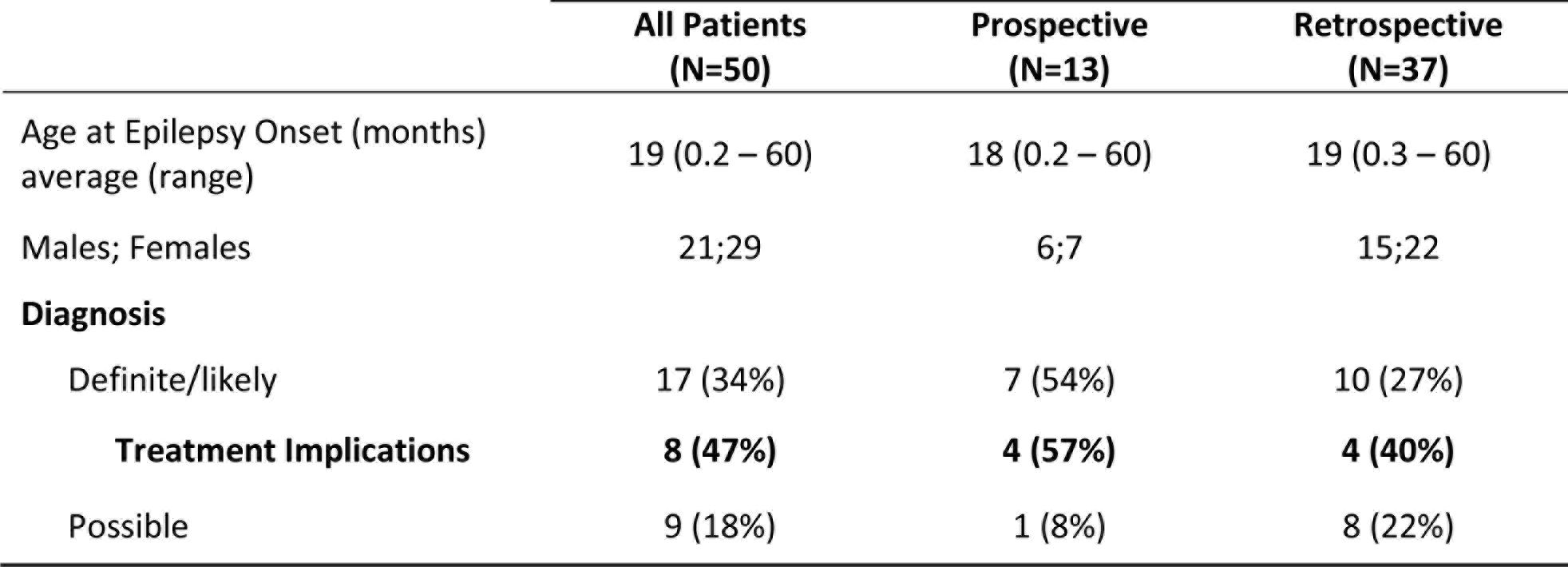
Results: Demographics and Diagnostic Yield

|  | **All Patients**<br>**(N=50)** | **Prospective**<br>**(N=13)** | **Retrospective**<br>**(N=37)** |
| --- | --- | --- | --- |
| Age at Epilepsy Onset (months)<br>average (range) | 19 (0.2 – 60) | 18 (0.2 – 60) | 19 (0.3 – 60) |
| Males; Females | 21;29 | 6;7 | 15;22 |
| **Diagnosis** |  |  |  |
| Definite/likely | 17 (34%) | 7 (54%) | 10 (27%) |
| **Treatment Implications** | **8 (47%)** | **4 (57%)** | **4 (40%)** |
| Possible | 9 (18%) | 1 (8%) | 8 (22%) |

### Diagnostic Yield

A definite or likely diagnosis was made in 17/50 patients (34%). A possible diagnosis was identified in another 9 (18%)(Supplementary Table 3). Eight of 17 patients (47%) were given a definite or likely diagnosis with potential treatment implications (Table 1). Pathogenic variants were identified in 15 genes and the majority were the result of *de-novo* mutations (12/17). The diagnostic yield was higher in the prospective (54%) than retrospective group (27%). Patients in whom a diagnosis was made had earlier onset epilepsy (mean 8.6 vs 27 months, t-test p-value<0.001), and global developmental delay and/or intellectual disability were more common (Table 3). Of 27 patients with epileptic encephalopathy a definite or likely pathogenic variant was identified in 12 (44%)(Supplementary Table 4).

### Treatment implications

A genetic disorder with specific therapeutic implications was diagnosed in eight patients (4 prospective and 4 retrospective). Clinical information, treatment changes, and impact are summarized (Table 2). Variants in *SCN5A*, incidental to patient phenotype but with treatment implications, were identified in 2 individuals (001, 067)(Supplementary Table 3). *SCN5A* mutations are implicated in cardiac arrhythmias with sudden death and, rarely, epilepsy (OMIM 300163). Both patients were evaluated by a Cardiologist and no abnormalities were found.

**Table 2.**
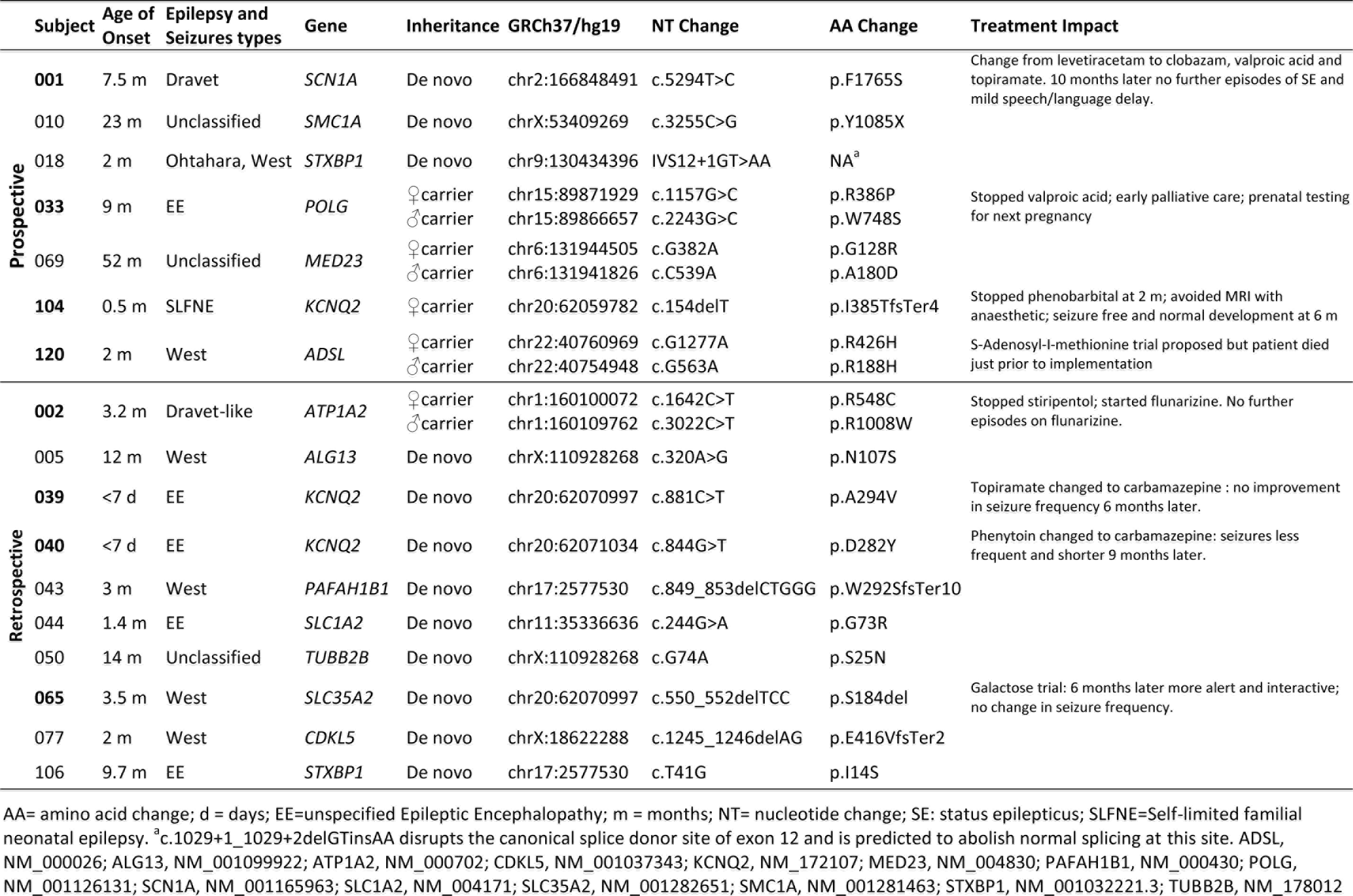
Patients with definite/likely diagnosis and treatment impact.

**Table 3.**
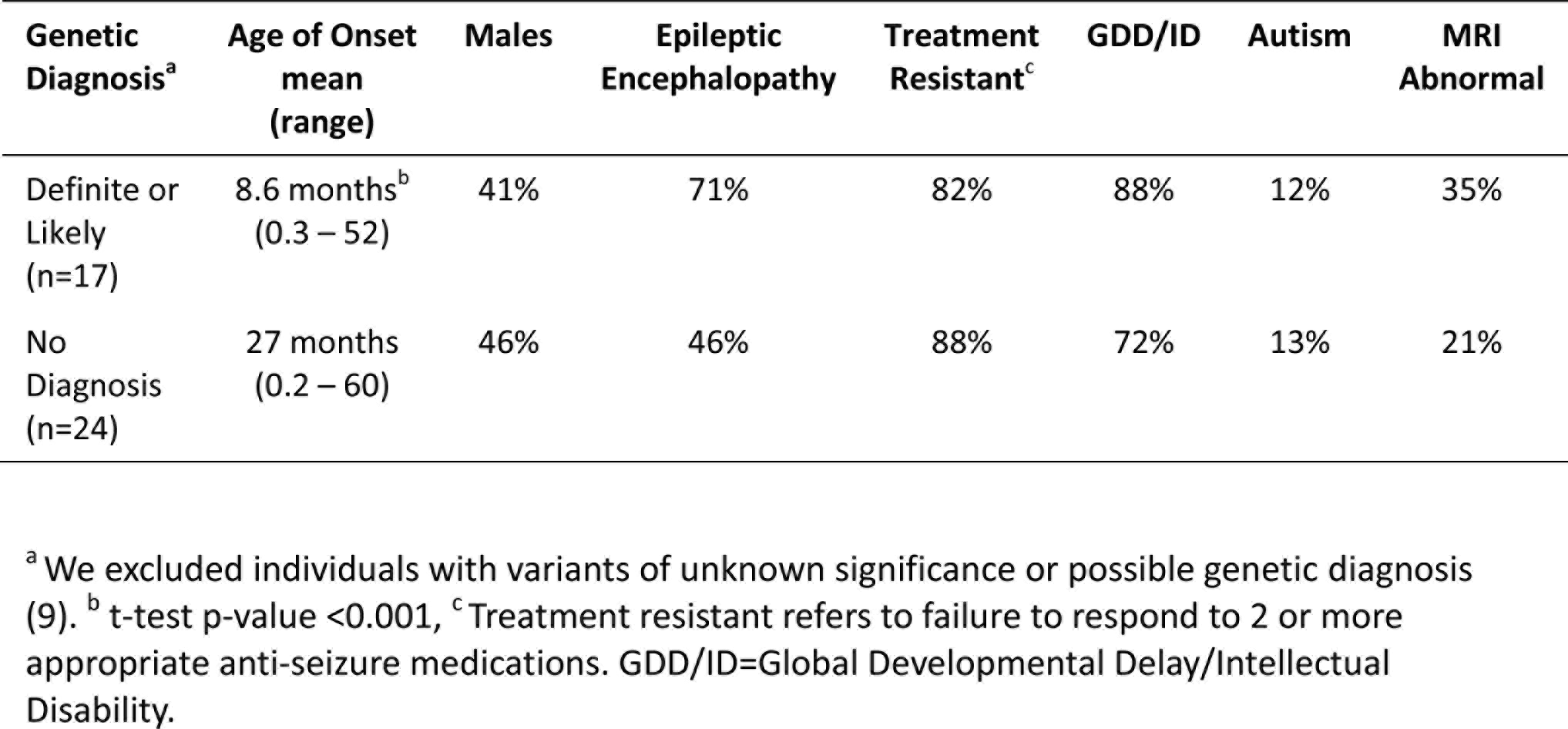
Clinical Features in Patients with and without a Genetic Diagnosis

| **Genetic Diagnosis<sup>a</sup>** | **Age of Onset mean (range)** | **Males** | **Epileptic Encephalopathy** | **Treatment Resistant<sup>c</sup>** | **GDD/ID** | **Autism** | **MRI Abnormal** |
| --- | --- | --- | --- | --- | --- | --- | --- |
| Definite or Likely (n=17) | 8.6 months<sup>b</sup><br>(0.3 – 52) | 41% | 71% | 82% | 88% | 12% | 35% |
| No Diagnosis (n=24) | 27 months<br>(0.2 – 60) | 46% | 46% | 88% | 72% | 13% | 21% |

<sup>a</sup> We excluded individuals with variants of unknown significance or possible genetic diagnosis (9). <sup>b</sup> t-test p-value <0.001, <sup>c</sup> Treatment resistant refers to failure to respond to 2 or more appropriate anti-seizure medications. GDD/ID=Global Developmental Delay/Intellectual Disability.

### Comparative time to diagnosis

The mean time to genetic diagnosis, from study enrolment with genetic counselling to research validation of the variant was 38 days (20-70) for the prospective group, 48 days (26-105) for the retrospective group, and 44 days (21-105) overall. The mean time from epilepsy diagnosis to research validation of genetic diagnosis was 143 days (42-242) for the prospective group, and 2,172 days (42-6,040) or ~6 years for the retrospective group.

### Cost Analysis

Point estimates and 95% confidence intervals, based on bootstrapped standard errors (1000 times with replacement) for each category of diagnostic test by cohorts, were calculated (Table 4). All cost estimates use rates effective for the 2014-2015 fiscal year. The mean total cost related to the diagnosis of epilepsy was $4,524 (range $1,223-$7,852) for the prospective cohort and $8,344 (range $3,319-$17,579) for the retrospective cohort. Diagnostic imaging and electrophysiological tests comprise >60% of total epilepsy-related diagnostic costs. The mean for diagnostic imaging testing constituted $1,391 and $3,276, for prospective and retrospective cohorts, respectively. The mean for electrophysiological testing constituted $1,353 and $2,731, for prospective and retrospective cohorts, respectively.

**Table 4.** Average diagnostic investigation cost per patient

|  | Mean | Bootstrap<br>Std. Err. | 95% CI |  | Range |  |
| --- | --- | --- | --- | --- | --- | --- |
|  |  |  |  |  | Min | Max |
| **Combined Diagnostic cost** |  |  |  |  |  |  |
| Retrospective | \$8344.27 | \$556.97 | \$7252.61 | \$9435.90 | \$3318.50 | \$17578.8 |
| Prospective | \$4524.07 | \$497.57 | \$3548.85 | \$5499.29 | \$1223.18 | \$7852 |
| **Lab tests** |  |  |  |  |  |  |
| Retrospective | \$1333.5 | \$83.71 | \$1169.42 | \$1497.56 | \$123.30 | \$3129.91 |
| Prospective | \$1151.32 | \$135.14 | \$886.43 | \$1416.19 | \$209.75 | \$1959.19 |
| **Genetic tests** |  |  |  |  |  |  |
| Retrospective | \$1179.55 | \$98.80 | \$985.90 | \$1373.20 | 0 | \$2279.34 |
| Prospective | \$633.187 | \$144.27 | \$350.41 | \$915.96 | 0 | \$1720 |
| **Diagnostic Imaging** |  |  |  |  |  |  |
| Retrospective | \$3276.10 | \$214.10 | \$2856.47 | \$3695.74 | \$1460 | \$6836 |
| Prospective | \$1391.38 | \$150.26 | \$1096.86 | \$1686 | \$630 | \$2290 |
| **Electrophysiological** |  |  |  |  |  |  |
| Retrospective | \$2731.22 | \$376.26 | \$1993.75 | \$3468.70 | 0 | \$8460.45 |
| Prospective | \$1353.12 | \$315.64 | \$734.4 | \$1971.78 | \$188.01 | \$3572.19 |

Our alternative scenario for diagnostic testing is MRI, EEG, chromosome microarray (CMA) and WES testing with Sanger sequencing validation, which amounts to $3,234 per patient (Supplementary Table 5). The difference in mean total cost related to the diagnosis of epilepsy for prospective ($4,524) and retrospective ($8,344) groups, exceeds the cost of our diagnostic alternative ($3,234). The potential average savings of targeted WES in the diagnostic workup constitute $1,290 per prospective patient and $5,110 per retrospective patient.

## Interpretation

Studies have supported high-throughput panel sequencing as a first-tier testing approach over similarly targeted WES for several diseases based on diagnostic yield, coverage, and cost-savings (7,28). However, a recent comparative coverage analysis limited to disease-causing variants identified through panels demonstrated that targeted WES detects ≥98.5% of those mutations (29), and that both approaches have comparable diagnostic yield. A major advantage of WES over panels is the ability to sequence the entire coding genome. Such comprehensive assessment can facilitate re-analysis for novel genes as they are implicated (in the course of this study ~7 genes were identified in seizure disorders and could be examined). Panel sequencing cannot include such contemporary targets.

The clinical utility of targeted WES with Sanger validation (limited ≤3 variants/exome) is supported by the identification of a definite or likely diagnosis in 17/50 (34%) patients and a possible diagnosis in an additional 9/50 (18%)(Table 1). A higher yield was found in the prospective group with new-onset epilepsy and supports earlier testing, though the number of patients is small. The retrospective group had already undergone extensive clinical testing that was non-diagnostic. Nevertheless, our ability to still identify a genetic diagnosis supports the technology’s superior resolution, while related data on phenotypes, management and outcomes may yet inform clinical practice.

The diagnostic yield in our study is comparable to previous findings (2–6,30). Most variants were *de-novo* and the genetic causes identified were heterogeneous, with recurrent variants only identified in *KCNQ2* (Table 2). In a comparable cohort, positive results were identified by WES in 112/293 (38.2%) epilepsy patients (30). We concur that the diagnostic yield is likely affected by the characteristics of the group studied, sample size, platform used (gene panel or WES) and the timing of the study, given ongoing gene discoveries in epilepsy. In our study, patients with a genetic diagnosis were younger and more likely to have global developmental delay/intellectual disability compared to patients in which no genetic cause was found. Similar to a prior study (30), our patients with epileptic encephalopathies had a high rate of positive findings (44%).

Our results support the feasibility of targeted WES to rapidly provide clinically-confirmed genetic diagnoses in early-onset epilepsy. Time to Sanger sequencing validation from enrollment averaged 6 weeks which is similar to the 6-8 week turn-around-time quoted by most commercial testing labs. However, this estimate did not include the additional time required to obtain provincial government approval, on a case-by-case basis, to fund WES.

A timely genetic diagnosis is important when considering the potential for treatment impact and optimization of patient outcomes. For the seventeen patients with a genetic diagnosis, eight (47%) were identified to have a disorder with specific treatment implications; for all eight patients, an immediate change in medical management was made (Table 2). The number of genetic disorders identified to have specific treatments implications is likely to grow with ongoing advances in precision medicine.

In British Columbia, the average savings are estimated to be between $1,290 and $5,110 per patient, depending on whether they are new prospective referrals or retrospective. Of note, price estimates reflect academic and/or hospital costs rather than commercial costs, which are 1-5X higher. The Canadian sequencing costs cited are comparable to previous reports but will decrease as even higher throughput sequencing technologies become accessible (7–10). Current healthcare cost estimates are also conservative as patients without a genetic diagnosis will undoubtedly require additional clinic visits and inpatient hospital stays, including epilepsy monitoring unit admissions related to finding the cause of their condition. Of note, a targeted WES approach did not lead to a substantial increase in referrals for incidental findings. Overall, our findings show targeted WES may provide an effective end to an otherwise invasive, time consuming and costly diagnostic odyssey, with societal and economic benefits. Our results also support WES implementation beyond early-onset epileptic encephalopathies, as we have examined a larger and more diverse group of children (10).

### Limitations and strengths

Our study has several limitations, including small sample size although our diagnostic yield is comparable to previous studies (30). Incomplete coverage of the 565 genes analyzed was partially addressed as outlined in the methods. Proband-parent trio-based WES analyses were not used primarily for financial reasons. Analysis was restricted to 565 epilepsy genes, rather than the entire exome, to identify a genetic diagnosis as quickly as possible and to minimize secondary findings. Assessing relevance of secondary findings and proving pathogenicity of variants in novel candidate epilepsy genes is costly; thus, this approach was taken to maximize patient care and minimize cost. WES data from patients with initial negative results continues to be periodically reviewed for variants in newly described epilepsy genes. In subsequent WES trio analysis, a subset of families has helped identify novel genetic etiologies (31).

Although almost half of the diagnoses had treatment implications, the long-term impact on clinical outcome following genetically-informed therapeutic interventions is unknown. Early diagnosis and early intervention are important, but advances in precision medicine are also required.

The methods employed for cost analysis cannot replace a prospective randomized controlled trial (RCT) and may not have accurately assessed or included all healthcare costs related to an epilepsy diagnosis. However, an RCT assessing the effect of WES testing on healthcare costs is not yet a practical consideration. Our estimates are not a perfect or a complete description of the current diagnostic work-up, as test records are scattered across different electronic health records systems and paper charts. Data collation within an accessible unified health electronic record would help identify where additional savings are possible. In this study, indirect costs, and the psychosocial impact on the child and family were not measured.

### Conclusion/Summary

Targeted WES with limited Sanger sequencing validation is a rapid and minimally invasive test with potential to save costs within the Canadian healthcare system. An early genetic diagnosis may improve a patient’s clinical outcome and quality of life. Further research on larger cohorts is warranted to inform diagnosis, clinical outcome and precision medicine. Acknowledging the limitations of our study, targeted WES with Sanger sequencing validation substantially improves current practice and is recommended as the dominant diagnostic strategy.

## Acknowledgements

We thank the children and families that took part in this study. We gratefully acknowledge Dr. Suzanne Vercauteren MD, PhD (Director), Dr. William Gibson MD, PhD (Chair and Biospecimen Advisory Committee member), Tamsin Tarling (Biobank Administrative Manager) and the technical expertise of Katelin Townsend of the BC Children’s Hospital BioBank, which is supported by Mining for Miracles through the BC Children’s Hospital Foundation. We also thank the BC Children’s Hospital Department of Pathology & Laboratory Medicine and the BC Children’s Hospital EEG (Electroencephalogram) Department. We are also grateful to Dr Vesna Popovska MD, Katie Pezarro, and Giselle Hunt BSc, from the Neurology Research Team at BC Children’s Hospital; and Dr Hilary Vallance MD, and Dr Graham Sinclair PhD, from the Department of Pathology, Biochemical Genetics Laboratory at BC Children’s Hospital, for their assistance on this project.

## Contributors

Michelle Demos contributed to conception and design of the study; analysis and interpretation of data; review of patients clinically, obtaining funding; drafting the manuscript. Ilaria Guella contributed to the analysis and interpretation of data and revised the manuscript. Marna B. McKenzie executed the genetic study and contributed to the acquisition and analysis of genetic data. Cyrus Boelman contributed to clinical assessments; data acquisition and interpretation and revised the manuscript. Daniel M. Evans performed bioinformatics and data analysis. Sarah Buerki designed the REDCap database, contributed to data acquisition, analysis and interpretation of data, and revised the manuscript. Eric Toyota contributed to data acquisition. Linda Huh contributed to clinical assessments; data acquisition; and revising manuscript. Anita Datta, Aspasia Michoulas, Kathryn Selby Bruce Bjornson contributed to clinical assessments and data acquisition. Gabriella Horvath and Ramona Salvarinova performed biochemical clinical assessments and contributed to the interpretation of genetic data. Elena Lopez performed the clinical assessments and contributed to the interpretation of genetic data. Clara DM van Karnebeek contributed to the study design and variant interpretation. Erin Slade assisted preparation of execution of study. Patrice Eydoux performed cytogenetic assessment and interpretation of genetic data. Shelin Adam performed the genetic counselling, and contributed to data acquisition, review of literature, and revising manuscript. Margot I Van Allen contributed to genetic counselling, review of results and literature and revising manuscript. Tanya N Nelson contributed to analysis and interpretation of data, performed the clinical variant interpretation, revised the manuscript, and negotiated funding for variant clinical validation. Corneliu Bolbocean performed economics data acquisition, analysis and interpretation; drafted economics section of manuscript; and revised the manuscript. Mary B Connolly performed electroclinical phenotyping of patients; revised the manuscript and obtained funding. Matthew J Farrer contributed to conception and design of the study and to genetic analysis and interpretation of data; obtained funding for exome sequencing and Sanger validation, revised the manuscript. All of the authors approved the final version to be published and agreed to be accountable for all aspects of the work.

